# Covary: A translation-aware framework for alignment-free phylogenetics using machine learning

**DOI:** 10.1101/2025.11.13.687960

**Authors:** Marvin De los Santos

## Abstract

In large-scale phylogenetic analysis, incorporating translation awareness is critical to account for the genotypic and phenotypic dimensions underlying biological diversification. Covary is a machine learning-based framework that analyzes, clusters, and compares genetic sequences through alignment-free, translation-aware embeddings. By integrating codon-boundary and intra-sequence positional information into a unified vector representation, Covary encodes mutational patterns alongside translation-level constraints. This design enables discrimination of frameshift-inducing mutations, substitutions, and other biologically meaningful sequence variations relevant to evolutionary relationships. Despite inherent sensitivity to *k-mer*-based distortions, Covary accurately clustered sequences, identified species, and reconstructed phylogenetic trees across diverse datasets, including human TP53 variants, ribosomal gene markers (18S and 16S), and complete genomes from viral, bacterial, and archaeal taxa. The resulting topologies were comparable to those produced by multiple sequence alignment (MASA)-based implementations like ETE3, with near-linear scalability demonstrated by the successful analysis of nearly a thousand SARS-CoV-2 genomes within minutes. The versatility and interpretability of Covary across mutation-, gene-, and genome-level analyses underscore its potential as a biologically informed, data-driven tool for bioinformatics, comparative genomics, taxonomy, ecology, and evolutionary studies. Covary is available online at https://github.com/mahvin92/Covary or at https://covary.chordexbio.com.

## INTRODUCTION

Our knowledge of the complex relationships among life on Earth remains incomplete, if not, imperfect. Classifying, studying, and reconstructing these relationships continue to be error-prone, time-consuming, and resource-intensive tasks for biologists. Despite recent advances in computational biology, phylogenetic inference still hinges on assumptions and models that often oversimplify or misrepresent the selective and stochastic processes shaping molecular evolution (Dimayacyac et al., 2023; Som, 2015; Fleming et al., 2023).

Molecular phylogenetics has long relied on DNA or protein sequences as the primary basis for reconstructing relatedness (Kapli et al., 2023; Gupta and Vadde, 2023). While DNA sequences provide high-resolution information on mutational events, protein sequences capture the functional consequences of these changes through the genetic code (Dimayacyac et al., 2023; Radványi and Kun, 2021). The choice between nucleotide or amino acid for phylogenetic analysis remains a longstanding debate, as each approach offers distinct advantages and limitations (Kapli et al., 2023; Puente-Lelievre et al., 2025; Foster and Hickey, 1999). Regardless of approach, a shared consensus in modern phylogenetics is that the major bottlenecks to accurate and objective inference lie in three major areas: 1) the availability of robust statistical models, 2) the type and context of molecular data, and 3) the computational efficiency of available analytical methods (Gupta and Vadde, 2023; Kapli et al., 2023; Dimayacyac et al., 2023; Fleming et al., 2023).

Recent developments suggest that improved results depend on more effective use of the rapidly growing biological data across the tree of life (Piñeiro and Pichel, 2024; Gardner et al., 2025; Haag et al., 2022; Young and Gillung, 2020). Machine learning and alignment-free models have emerged as promising approaches to overcome the limitations of multiple sequence alignment (MSA), allowing flexible representations of molecular data that can generalize across scales of divergence (Zuo and Hao, 2015; Deng et al., 2025). However, most of these methods remain limited to compositional features, overlooking one of the most fundamental biological processes connecting genotype to phenotype, which is translation.

To date, translation awareness in machine learning-based molecular phylogenetics remains largely unexplored. Comparable approaches, such as codon-based models, have been shown to yield higher efficiency and greater power in estimating relatedness under substitution assumptions, owing to their richer parametric vocabulary within codon space (Gupta and Vadde, 2023). Such models are particularly advantageous in cases of high genetic degeneracy, where synonymous substitutions carry functional implications (e.g., codon usage bias). However, their computational burden often limits scalability, particularly when applied to deep phylogenies or non-protein-coding datasets (Miller et al., 2020; Kapli et al., 2023; Gupta and Vadde, 2023; Shapiro et al., 2006). Thus, there is a persistent need for a more computationally-optimized contextual representation of genetic data that transcends mere compositional or structural features. To advance molecular phylogenetics, it is imperative to integrate translation awareness to faithfully consider both the mutational and functional dimensions that jointly drive evolution.

In this study, Covary, an alignment-free and translation-aware encoding framework for phylogenetic analysis built on TIPs-VF (De los Santos, 2025), was evaluated across diverse data types and biological questions. Specifically, the objectives were to: (1) confirm that Covary retains alignment-free comparability with its predecessor, while introducing translation awareness; (2) assess whether translation-aware representations enhance discrimination of reading-frame-altering mutations; (3) demonstrate the capability of Covary for phylogenetic reconstruction using both marker genes and complete genomes; (4) evaluate its power for species identification and taxonomic resolution; and (5) gain insights about its applicability for phylogenomics. This paper describes the preliminary success of Covary, its limitations, and future directions.

## METHODS

### Data acquisition and processing

The complete genome sequences of SARS-CoV-2 variants KP.3.1.1, JN.1, XEC, MV.1, LP.8.1, NB.1.8.1, XDV.1, LF.7, and XEG were retrieved from The NCBI Viral Genomes Resource (Brister et al., 2015). The accession numbers of the sequences and their corresponding Pango lineages are listed in Supplementary Table 1 and their sampling/surveillance detection details can be retrieved from the source NCBI database. A total of 906 whole genome sequences were used. All sequences were pre-processed to start relatively, as the reference Wuhan-Hu 1 complete genome sequence (NC_045512.2), using the Seed Aligner implementation (https://github.com/mahvin92/Seed-Aligner).

The TP53 mutational profiles of GDC-TCGA Head and Neck Cancer (HNSC) cohort were retrieved using the UCSC Xena Browser (https://xenabrowser.net/). A total of 220 patient data with TP53 mutations and wildtype profiles were included in the analysis. The locations of mutation in the TP53 gene were determined by normalizing the chromosomal location of the affected nucleotides in the patients’ molecular profiles with the chromosomal location of the TP53 gene (chr17: 7,668,421..7,687,490) from the GRCh38 assembly. The derived locations were used to recreate the gene-level mutational profiles of the patients using the reference sequence NC_000017.11 in the Mutagen PX implementation (https://github.com/mahvin92/Mutagen-PX). This type of ‘simulated’ mutation generation was performed due to speed and ease of use, rather than going through the large and complex individual sequencing data of all patients included in the analysis. This reverse simulation was expected to yield similar sequences, since all genome assemblies and annotations utilized the same logic and workflow implemented in the analysis. Codon-bias, which may not be captured in this simulation was irrelevant in the analysis. The TCGA sample IDs of the cohort were listed in Supplementary Table 1.

Mammalian 18s rRNA (n=7) and bacterial 16s rRNA (n=607) gene sequences were retrieved from NCBI Nucleotide (https://www.ncbi.nlm.nih.gov/nuccore/). Unaligned 18S rRNA sequences of *Rattus norvegicus, Mus musculus, Bos taurus, Gorilla gorilla, Pan troglodytes, Homo sapiens*, and *Aspergillus niger* were analyzed directly using Covary and ETE3 (Huerta-Cepas et al., 2016). Randomly selected 16S rRNA sequences from bacterial genera including *Thermus, Mycoplasma, Escherichia, Staphylococcus, Pseudomonas*, and *Lactobacillus* were also used without preprocessing. The corresponding accession numbers, species names, and GenBank titles are provided in Supplementary Table 1.

Whole genome sequences from selected Archaea (*Methanosarcina acetivorans, Sulfolobus acidocaldarius, Methanobrevibacter smithii, Metallosphaera sedula, Thermococcus kodakarensis*) and Bacteria (*Stenotrophomonas maltophilia, Klebsiella variicola, Helicobacter pylori, Bacillus licheniformis, Neisseria meningitidis*) were obtained from the NCBI Genomes Database (https://www.ncbi.nlm.nih.gov/datasets/genome/). Additionally, 40 randomly selected viral genomes (based on top search results) were analyzed separately for phylogenomic inference. All sequences were used directly without preprocessing. Accession numbers and GenBank titles are enumerated in Supplementary Table 1.

### Phylogenetic reconstruction with Covary

#### Access and versioning

Covary can be accessed through GitHub (https://github.com/mahvin92/Covary/) or through the website (https://covary.chordexbio.com).

#### Covary execution

All analyses in this report were performed using Covary v2.1 (more information about the different versions of Covary is available at https://github.com/mahvin92/Covary/releases). The Covary implementation was tested using Google Colab (free tier), utilizing 12.7 GB system RAM, 15.0 GB GPU RAM, and 112.6 GB disk storage. The Covary encoder version 2025-Q3 was used in all experiments (https://github.com/mahvin92/Covary-encoder). Since the Covary encoder was derived from a previously described genetic encoding logic, TIPs-VF (see Supplementary Figure 1), the comparable performance of Covary encoder and TIPs-VF was initially verified (data not shown). The machine learning implementation on Covary utilized Keras, which has been previously described (De los Santos, 2025), with some modifications. Analysis using Covary was performed using the default parameters.

**Figure 1.**
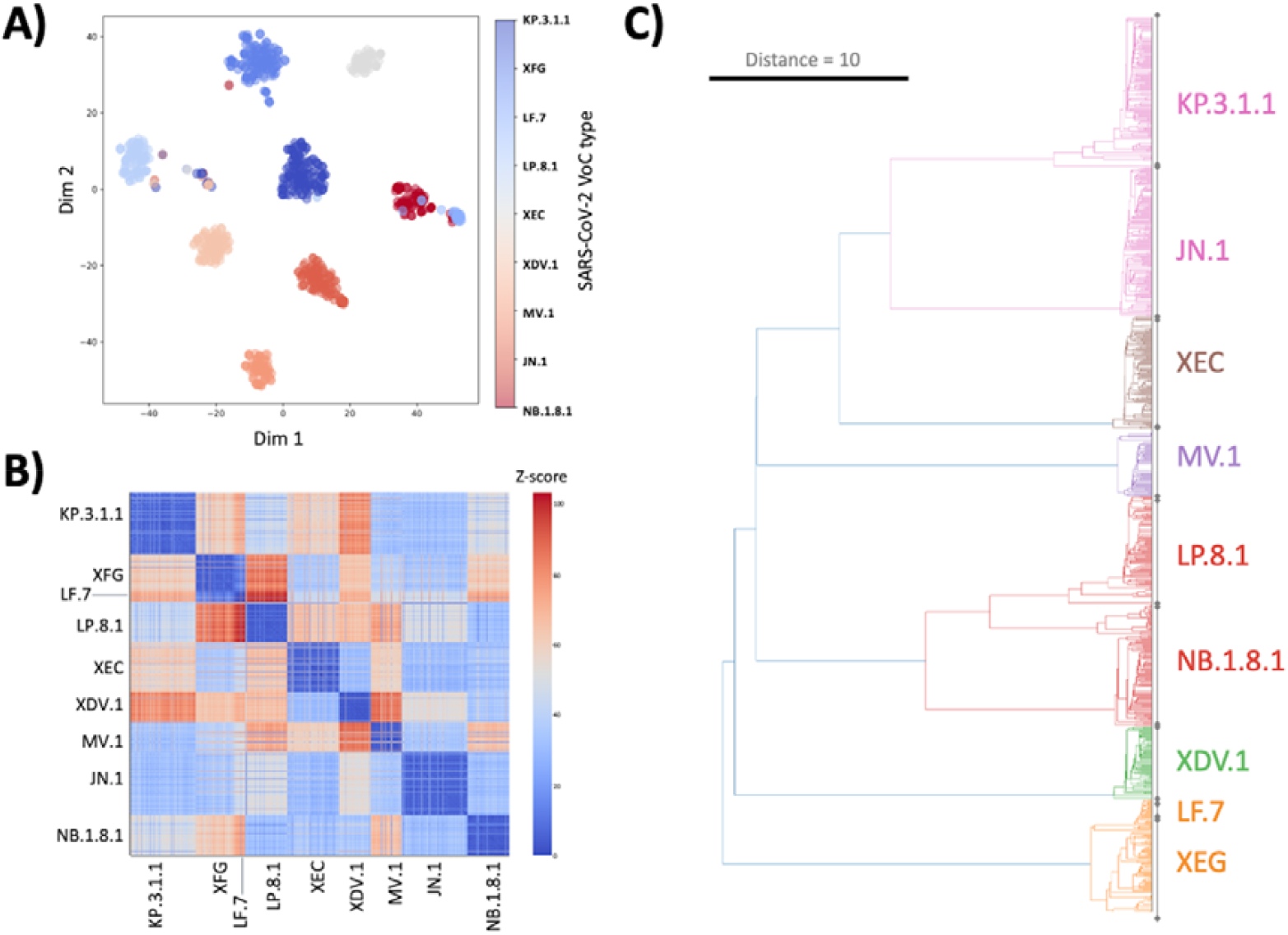
Clustering of whole genomes from different SARS-CoV-2 variants using Covary (n=906). A) Dimensional projection of the sequence embeddings generated by Covary using t-SNE. Clustering patterns were visually examined by assessing the homogeneity of color distribution within each cluster. Covary was able to discern and reclassified previously identified variants into different groups, an interesting foundational evidence to re-visit whole genome-based methods used in SARS-CoV-2 variant classification. B) Pairwise heat map representation of the vector distances for each genome sequence used to analyze the lineage-specific relationships of the Euclidean distance matrices derived from the t-SNE embeddings. C) Reconstructed phylogenetic tree based on the vector distances of each sequence. The tree was inferred using the t-SNE embeddings and constructed using single-linkage method. *Dim 1 = Dimension 1; Dim 2 = Dimension 2; Distance = branch length*.

#### Covary results

Covary infer phylogenetic relationship by interrogating the distance matrices of vector embeddings through clustering (vector projections using PCA, t-SNE, and UMAP), pairwise distance comparison (Euclidean method), and hierarchal clustering (by complete, single, average, and Ward methods). All these data were generated per each analysis and the most sensible prediction and possible model for phylogenetic estimation were selected by manual tree search, as described below.

#### Manual tree search

PCA, t-SNE, and UMAP create different reductional spaces, which introduce some form of variability in phylogenetic inferences. However, initial performance optimization of Covary showed that distance matrices derived from PCA were conducive for relationship estimation in sequences with very minimal differences (e.g., point mutations, single nucleotide polymorphism); whereas, t-SNE was found to assist in developing competitive models for deriving relationships in sequences with a minimal degree of complexity (e.g., internally gapped sequences), and UMAP supports estimation with large variability in sequence differences (e.g., fusion genes). For phylogenetic inference, t-SNE and PCA were used in manual tree search, unless specified in text or figure captions. Resulting topologies were inspected across all linkage methods, and the most plausible trees were selected based on visual fit and correspondence to reference trees generated by conventional tools such as ETE3.

#### Post-run analyses

Covary outputs include embedding coordinates (dim1, dim2), pairwise distance matrices, and hierarchical dendrogram linkage data, all compiled in the Covary report. Embedding and distance matrix data were subsequently subjected to statistical analysis.

### ETE3 implementation

Phylogenetic analysis was conducted using the ETE3 toolkit (Huerta-Cepas, et al., 2016) implemented in the GenomeNet platform (https://www.genome.jp/tools/ete/). The nucleotide sequences in FASTA format were uploaded directly to the web interface. Default settings were applied throughout the analysis. The Aligner was set to MAFFT for multiple sequence alignment, and IQ-TREE was used as the tree builder for phylogenetic inference.

### Statistical analysis

All statistical analyses, not executed in Covary, were performed using jStat.js (https://github.com/jstat/jstat) through a web-based implementation on PlethoCalc (https://plethocalc.plethoryt.com). Normality test was used to assess normal distribution of distance matrices (dim1 and dim2), descriptive statistics was used to assess differences of data clusters, and hypothesis test was used to assess differences in the observed clustering patterns. Specifically, one-way ANOVA was used to compare differences of groups with n > 2, while one-sample t-test was used to assess statistical significance in groups with heterogenous sample size of n ≥ 1. F-statistic, t-statistic, and p-values were recorded. P-values < 0.001 was considered statistically significant.

## RESULTS

### Alignment-free and translation-aware features of Covary

Covary uses a genetic representation module known as the Covary encoder, which was built using TIPs-VF (De los Santos, 2025). The Covary encoder extends TIPs-VF by using non-frequency associative *k-mer* units (default k = 6) with position awareness and explicitly encoding codon boundaries by concatenating codon-level representations for the three possible reading frames into a single unified representation. Unlike TIPs-VF, which produces three independent sequence products (one per frame), Covary encoder was enhanced to generate one compact codon-centric signature that preserves frame context while remaining alignment-free (Supplementary Figure 1). This design confers two practical advantages: 1) improved discrimination of variants that affect translation (frameshifts, in-frame indels, synonymous vs nonsynonymous changes), 2) and robustness to truncated/gapped sequence inputs where alignment is unreliable. To evaluate these properties, the clustering of dominantly circulating SARS-CoV-2 variants (as of August 2025, see METHODS) using whole-genome sequences and phylogenetic placement of TP53 mutant sequences from TCGA head and neck cancer (HNSC) patients were analyzed.

Using viral whole genomes sampled across their surveillance period (see METHODS for the list of sequence accession numbers), Covary representations were used to generate vector embeddings, compute pairwise distances, and perform hierarchical clustering. Genome sequences that correspond to the same variant class formed cohesive clusters in both the t-SNE projection and heat map pairwise cluster (Figure 1A–B). This observation contrast with previous findings where UMAP outperformed clustering projection by t-SNE and PCA in SARS-CoV-2 mutation dataset (Hozumi et al., 2021). It is hypothesized that the performances of these different dimensional reduction methods are influenced by the degree of *k-mer* disparity, stemming from variability in sequence differences (e.g., base mutations, sequences gaps, GC%, and compositional frequency). This principle has been previously demonstrated by Nanduri et al., 2024, which prompted the pre-optimization of Covary for different sequence anomalies such as single-base mutations, short indels, and gene fusions (see METHODS, Manual tree search).

Likewise, Covary was able to generate a representative phylogenetic tree of the variants while also inferring close relationship of sequences based on the expected groupings (Figure 1C). Cluster purity analysis by comparing the vector dimensional projections (dim1 and dim2) using normal distribution test and ANOVA showed that the observed clusters were unlikely to have formed randomly [F-statistics: 1: 717.60 (dim 1) and 1494.60 (dim 2), p <0.001 for both; Supplementary Figure 2]. These results indicate that Covary retained alignment-free discrimination at a whole-genome scale and grouped epidemiologically and genetically related SARS-CoV-2 lineages, at the genome scale, without explicit alignment steps.

**Figure 2.**
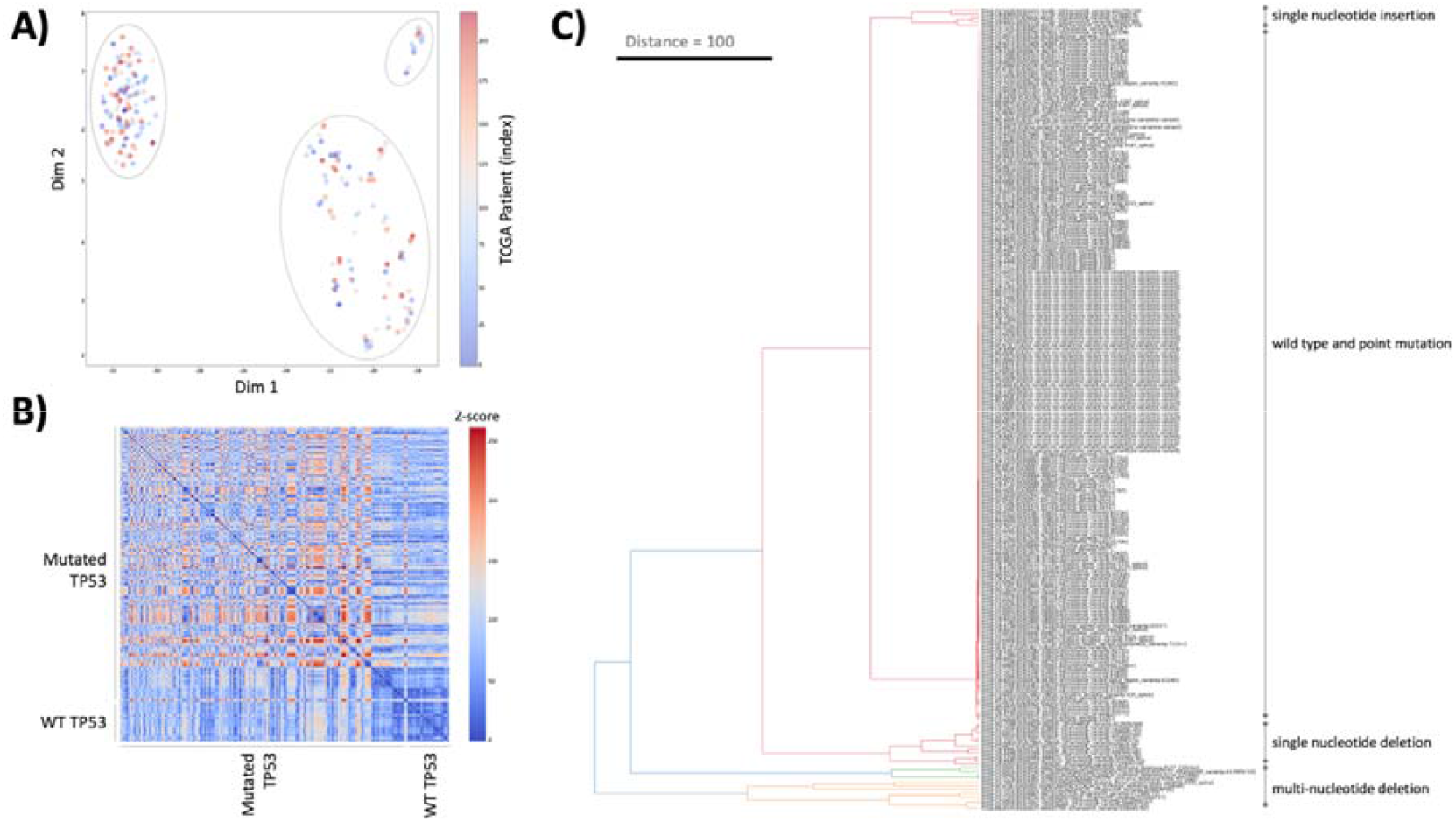
Demonstration of translation-awareness by Covary using the TP53 mutational variants from the GDC-TCGA HNSC cohort (n=220). A) Dimensional projection of the Covary-derived embeddings using PCA (principal component analysis). Clustering was visually examined by assessing the spatial proximity and separation of vector points across variant classes. B) Pairwise heat map representation of the vector distances among all TP53 sequences used to analyze mutational relationships of the different clinical variants. The distance matrices used were derived from the PCA embeddings. C) Reconstructed phylogenetic tree based on PCA-derived vector distances. The tree was inferred and generated using complete linkage method. WT = wild type; Dim 1 = Dimension 1; Dim 2 = Dimension 2; Distance = branch length.

Next, Covary was evaluated for its ability to distinguish sequence changes that alter translation (frameshifts, in-frame indels) from changes that do not (missense, synonymous substitutions), a key feature demonstrating its translation-awareness. TP53 sequences were reconstructed for HNC patients using GDC-TCGA mutation records and the Mutagen PX reconstruction pipeline: wildtype (WT) sequence was taken from the reference, mutated sequences were generated by applying the patient-level variant list to the WT (see METHODS).

Hierarchical clustering and heatmap of pairwise distance matrices partitioned the TP53 dataset into three probable groups: (i) sequences matching WT (non-variant), (ii) sequences harboring predominantly single-site substitutions (missense / synonymous), and (iii) sequences with indels that alter reading frames or create extensive local sequence disruption (Figure 2A). Covary effectively separated sequences with frameshift-causing indels from both WT and single-substitution sequences (Figure 2B). Interestingly, within the indel cluster, Covary further differentiated single-nucleotide insertions from single-nucleotide deletions and from more complex multi-nucleotide deletions, demonstrating sensitivity to the type of reading-frame and sequence length disruptions (Figure 2C). A normal distribution test and one-sample t-test on the Covary distance matrices suggested the non-random pattern in the observed TP53 mutational clusters [t-statistics: −65.38 (dim 1) and 51.35 (dim 2), p <0.001 for both; Supplementary Figure 3]. These analyses support that the codon-aware encoding of Covary captures translation-relevant signals not accessible to pure nucleotide composition methods.

**Figure 3.**
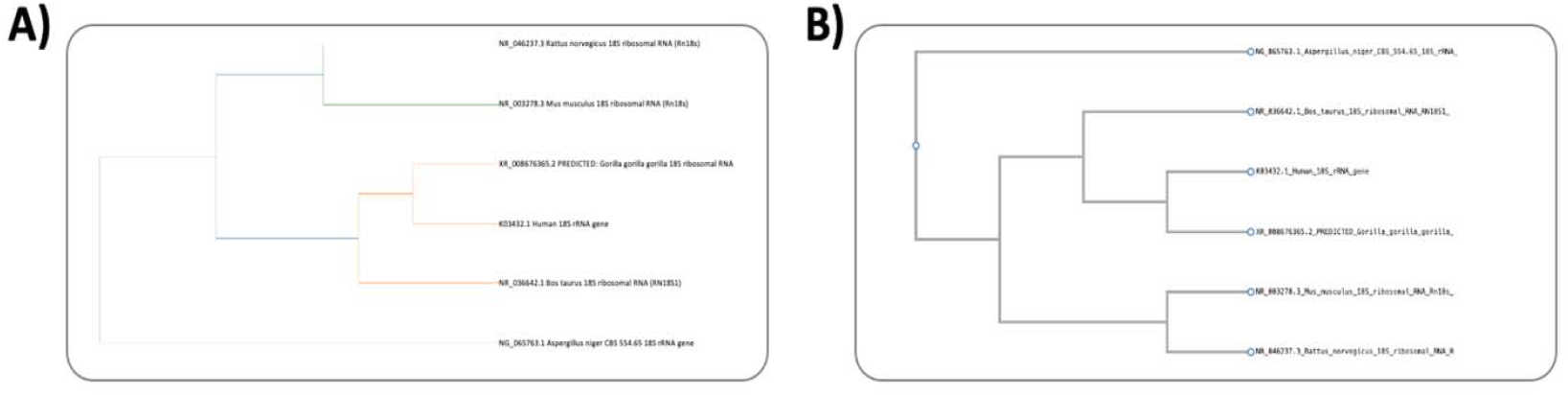
Representative gene tree reconstruction using 18s rRNA marker. A) Phylogenetic relationship inferred by Covary, with the tree reconstructed using complete-linkage clustering of the vector distances obtained from t-SNE embeddings. B) Reference phylogenetic tree generated by ETE3 using default parameters (MAFFT alignment and IQ-TREE inference). The 18S rRNA sequence of *Aspergillus niger* was designated as the outgroup, while sequences from mammalian representatives (*Rattus norvegicus, Mus musculus, Bos taurus, Gorilla gorilla, Pan troglodytes, Homo sapiens*) were treated as the ingroup. Tree topologies were similar between Covary and ETE3.

### Phylogenetic reconstruction using Covary

Covary was evaluated for its ability to produce biologically coherent gene trees. A benchmark 18S rRNA dataset representing six mammalian species (*Rattus norvegicus, Mus musculus, Bos taurus, Gorilla gorilla, Pan troglodytes, Homo sapiens*) with *Aspergillus niger* as an outgroup was analyzed. Covary inferred a topology that separates non-mammalian *A. niger* as the outgroup and partitions mammals into two major clades: rodents on one side and primates + ungulates on the other (Figure 3A). To validate topology consistency, the same tree was reconstructed using the ETE3 toolkit and observed concordant grouping, where *A. niger* sits as outgroup and primates and ungulates associating more closely with each other than either with rodents (Figure 3B). Branch length comparison was not performed due to the difference in the derivation of genetic changes between *k-mer*-based and MSA-based tools.

### Species identification with Covary

The ability of Covary to identify ingroup from outgroup sequences and to place ‘unknown-labelled’ sequences appropriately within a set of references was examined. Distance-based clustering in Covary correctly designated the viral genome as an outgroup relative to the bacterial 16S ingroup (Figure 4), recapitulating the same broad separation produced by alignment-based ETE3 analysis (Supplementary Figure 4). Further, when two unknowns were intentionally used as controls, SARS-CoV-2 isolate (PV95068.1) and a *Staphylococcus spp*. 16S (NR_036791.1), both Covary and ETE3 placed each unknown into the expected outgroup or ingroup clade, respectively. This suggests the similar application of Covary and MSA-based tools like ETE3 in species identification. Although comparing viral whole genomes to bacterial rRNA is not typical in phylogenetic analysis, this exercise demonstrates the robust separation across extreme sequence divergence by Covary. Contrary to assumptions that tools and frameworks for big data phylogenetics are susceptible to systematic errors and methodical incongruencies (Young and Gillung, 2020), Covary shows that phylogenomic tools can yield results that are consistent with traditional methods. Overall, the result recognizes the potential of Covary for broad phylogenetic workflow without explicit alignment or dependence on label assignment.

**Figure 4.**
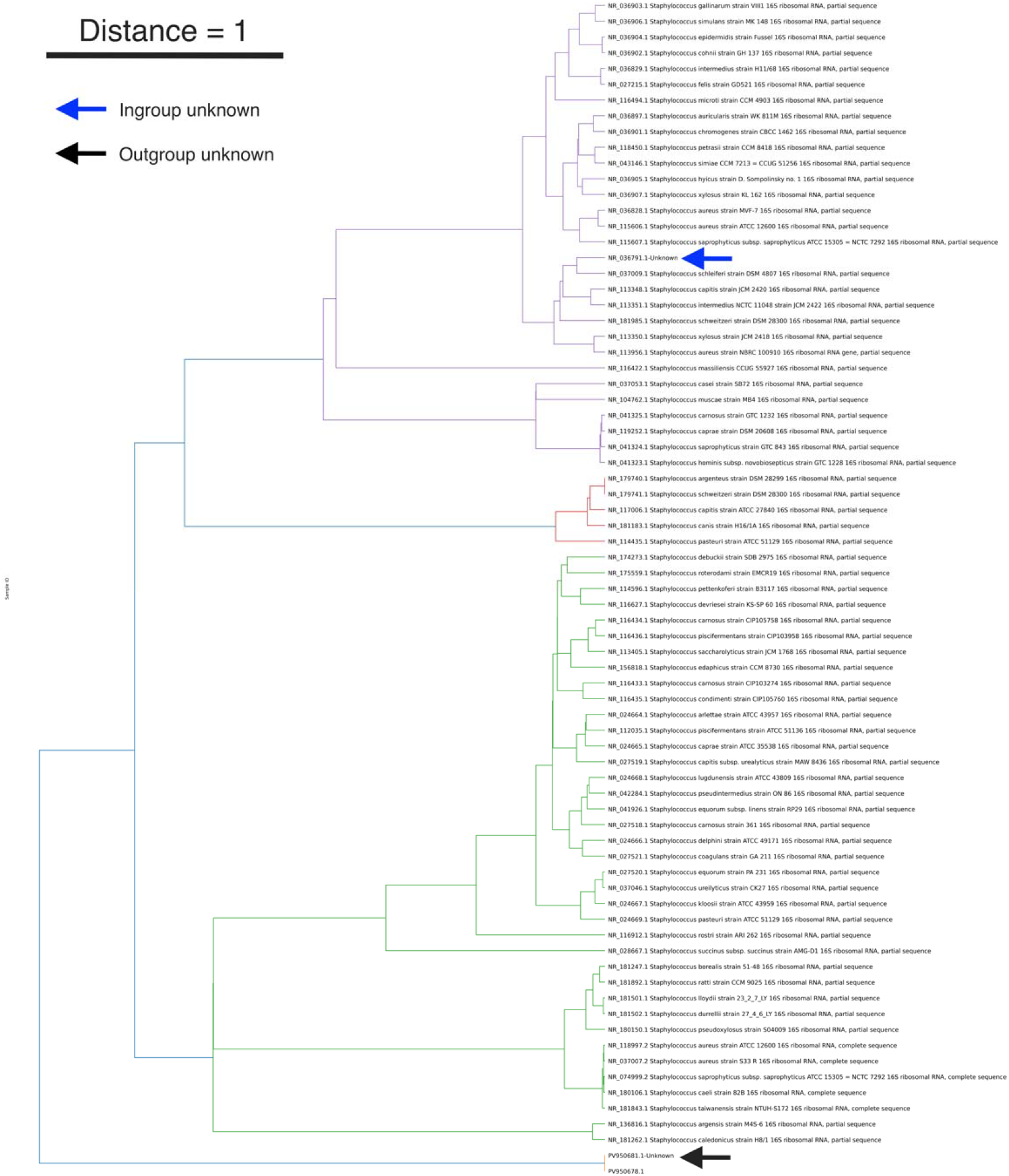
Illustrative workflow for species identification and outgroup detection using Covary (n=94). The translation-aware vector embeddings accurately grouped 16S rRNA sequences by genus (e.g., *Staphylococcus*), correctly clustered a taxonomy-verified *Staphylococcus* sequence of unknown label within its expected clade, and positioned viral outgroups (SARS-CoV-2 whole-genome sequences) at distinct boundaries, demonstrating accurate ingroup–outgroup discrimination.

### Resolving taxonomic order with Covary

To probe taxonomic resolution, Covary was applied to a curated multi-genus 16S rRNA dataset encompassing *Thermus, Mycoplasma, Escherichia, Staphylococcus, Pseudomonas*, and *Lactobacillus*. At the genus level, Covary grouped sequences into well-defined clades consistent with genus annotations (Figure 5). Notably, within *Lactobacillus*, Covary resolved closely related species pairs and in some cases distinguished strain/subspecies clusters that align with known intraspecific diversity (e.g., clustering of all sequences from subspecies or strains of *L. delbrueckii*; with similar fidelity for *L. kefiranofaciens*). Although Covary elevates translation-awareness in resolving genus-level inferences, these results indicate that its codon-boundary profiling can also be applied to sequences that do not encode proteins, such as 16S rRNA, indicating that the Covary framework could be adaptable to both protein-coding and noncoding structural RNA genes.

**Figure 5.**
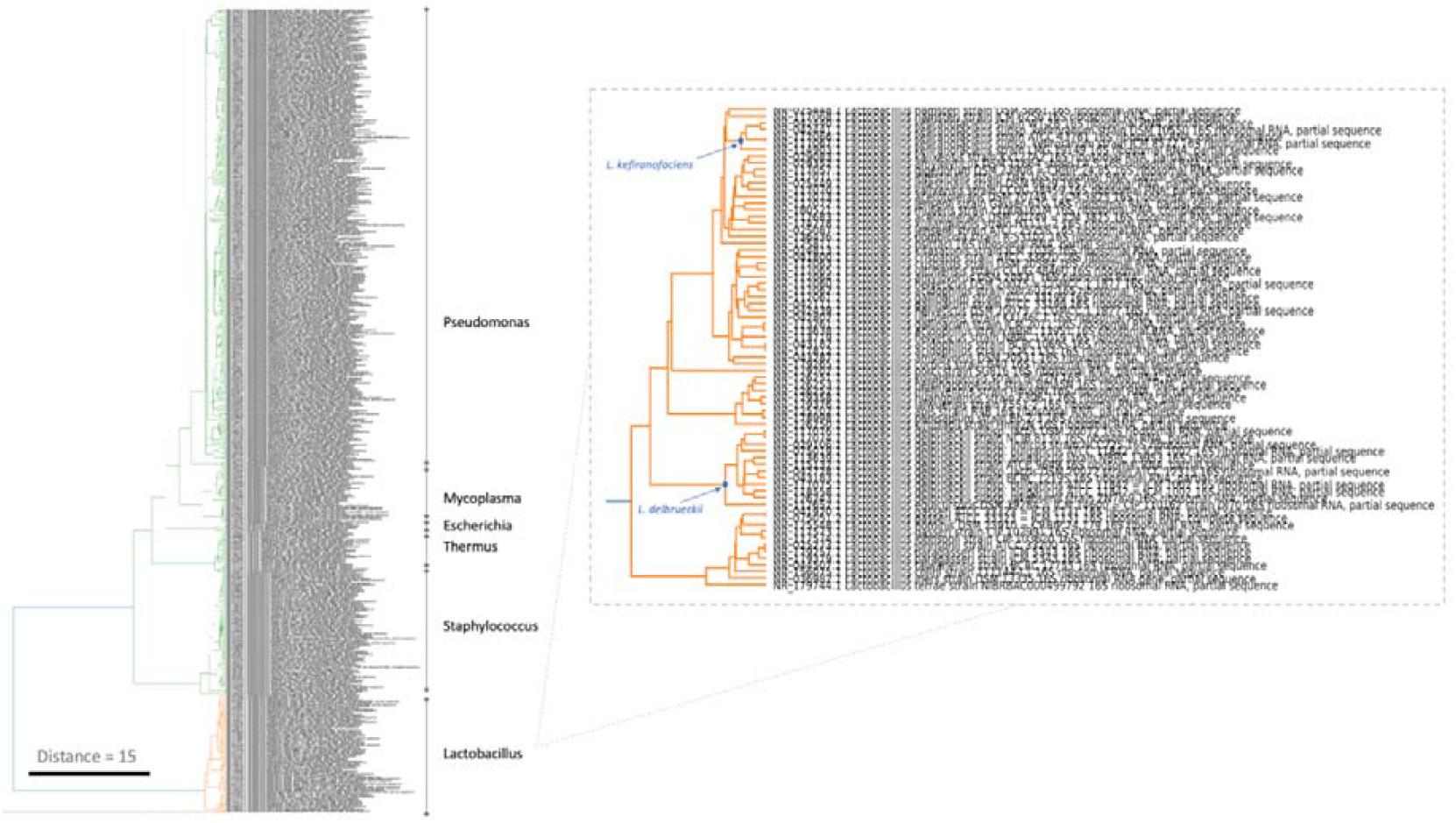
Taxonomic resolution across bacterial genera using Covary. Covary accurately clustered 16S rRNA sequences into genus-consistent clades (*Psuedomonas, Mycoplasma, Escherichia, Thermus, Staphylococcus, Lactobacillus*) and further distinguished closely related strain/subspecies, demonstrating its sensitivity in detecting taxonomic relationships even from non-protein-coding sequences. These findings indicate that Covary can be effectively integrated into conventional phylogenetic and taxonomic workflows.

### Phylogenomic analysis with Covary

The successful reconstruction of SARS-CoV-2 relationships using complete genome sequences, described above, provided an important proof of concept that an alignment-free, translation-aware approach could be effectively used for species-level phylogenetics. This observation motivated the broader application of Covary for phylogenomic inference across diverse viral taxa, with the goal of determining whether the same principles extend to complex datasets.

To evaluate this, Covary was tested on a random set of viral genomes, including complete genomes, coding sequences, and segmented genomes from a variety of virus families. The inclusion of such a heterogenous dataset that range from single-stranded RNA viruses to large double-stranded DNA viruses, was intended to assess the capacity of Covary to generalize across genome architectures and sequence lengths without requiring alignment or compositional normalization. Remarkably, Covary consistently clustered sequences according to known evolutionary relationships (Figure 6). Members of the *Coronaviridae* family, such as SARS coronavirus Tor2, Duck coronavirus DK/GD/27/2014, MERS-related coronavirus HCoV-EMC/2012, and Human betacoronavirus 2c EMC/2012, were grouped together, reflecting their shared ancestry and confirming that translation-aware embeddings can capture taxonomic coherence even among distantly related coronaviruses. Similar taxonomic fidelity was observed across other viral groups, including Influenza viruses, Monkeypox virus, and White spot syndrome virus, demonstrating that translation-aware representation by Covary retains sufficient biological signal for meaningful phylogenetic reconstruction across disparate viral families.

**Figure 6.**
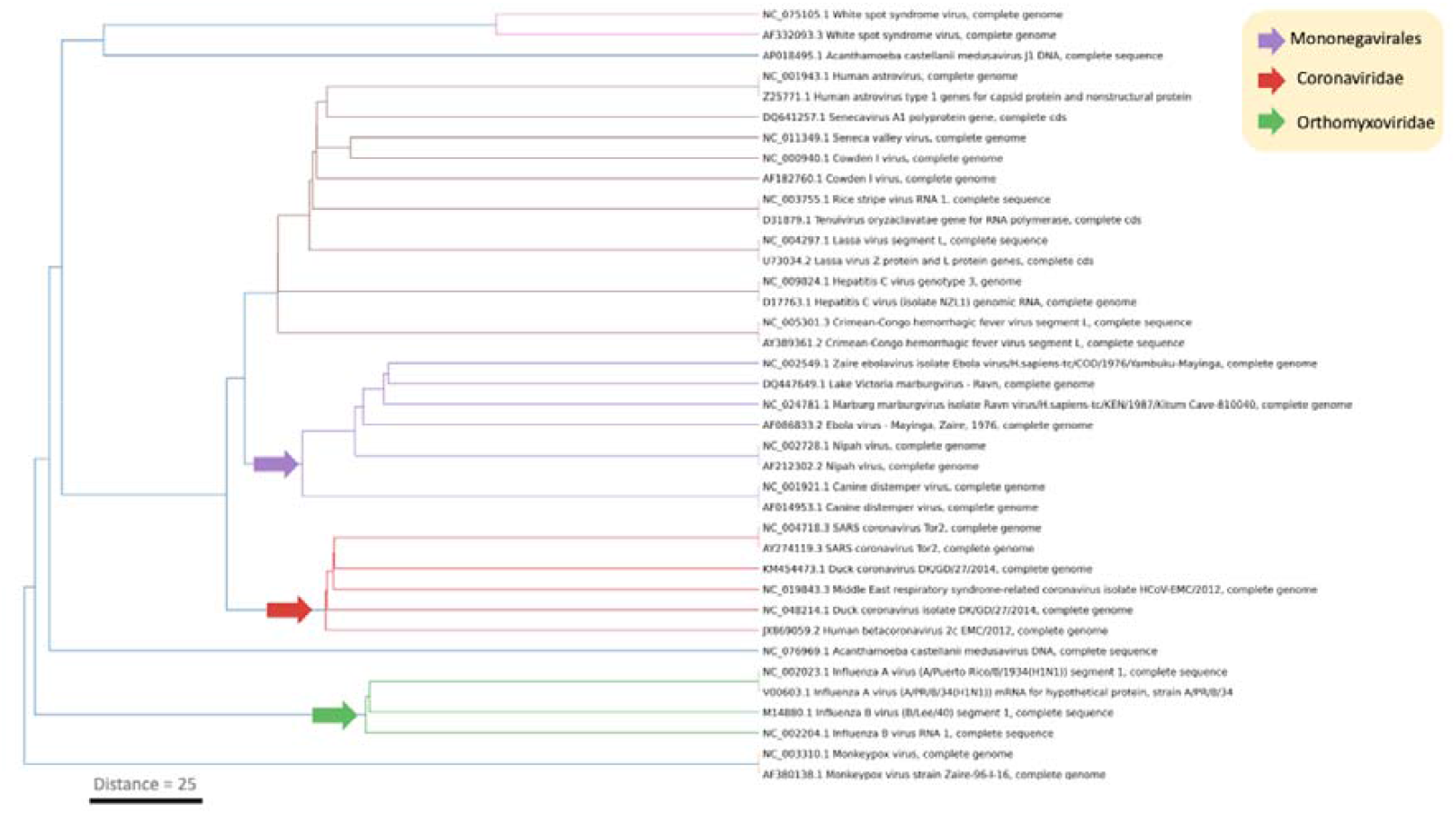
Phylogenomic application of Covary across diverse viral genomes. Covary correctly grouped viral genomes according to established taxonomic relationships, clustering related species and families such as *Coronaviridae, Filoviridae*, and *Paramyxoviridae*. The model also identified homologous regions among distantly related viral genomes, highlighting its ability to infer functional and evolutionary relationships directly from unaligned, translation-aware sequence embeddings.

From this random selection of viral species, a particularly striking observation emerged from the clustering of viruses belonging to the order *Mononegavirales*. Covary correctly positioned members of the *Filoviridae* (e.g., Ebola virus) and *Paramyxoviridae* (e.g., Nipah virus) families in close proximity, consistent with their known evolutionary relationship. This demonstrates that Covary not only distinguishes between major viral groups but also resolves fine-scale relationships between closely related families based on translation-aware sequence embeddings. Additionally, when the *Tenuivirus oryzaclavate* RNA polymerase gene coding sequence was analyzed alongside the genomic RNA 1 segment of Rice stripe virus, which encodes the same polymerase gene, Covary accurately clustered the two together. This result highlights that Covary identifies homology through intrinsic sequence features rather than relying on metadata or annotation, underscoring its data-driven ability to detect functional and evolutionary correspondence.

Covary was next applied to the more complex and larger genomes of Bacteria and Archaea to explore its scalability and sensitivity to deeply diverged phylogenomic signals. Analysis of complete archaeal and bacterial genomes using Covary revealed two major clades with unexpected inter-domain relationships (Supplementary Figure 5). Clade 1 gave rise to *Stenotrophomonas maltophilia*, whose ancestral lineage subsequently diverged into *Klebsiella variicola* and *Methanosarcina acetivorans*. Clade 2 split into two branch: one leading to *Methanobrevibacter smithii* and *Helicobacter pylori*, and the other showing successive splits among *Bacillus licheniformis, Metallosphaera sedula, Neisseria meningitidis*, and finally a split between *Thermococcus kodakarensis* and *Sulfolobus acidocaldarius*. These patterns are consistent with documented perturbations, rather than errors, in the contextual understanding of whole genome phylogenetics (Young and Gillung, 2020). Traditional assumptions of relatedness have been built on limited (‘minimum sequence’) datasets rather than whole genomes, and biologists are still in the process of deciphering and normalizing the process of whole genome-based phylogenetics (Yu et al., 2024; Zhou et al., 2021).

The result suggests that some archaeal species cluster closer to bacterial taxa than expected, likely reflecting a combination of horizontal gene transfer, genome-wide similarities, convergent evolution, variable evolutionary rates, and other methodical or compounding errors that are consistent with previous observations (Sakoparnig et al., 2021; Avni and Snir, 2020; Wang et al., 2024; Young and Gillung 2020). Regardless of these factors, this result suggests that despite the increase in genome size and structural complexity of data used for phylogenomic inference, Covary retained its capacity to organize sequences into representative clusters. This feature is a critical first step into fully realizing a system that interrogates whole genome data for faithful and more sensible phylogenetics reconstruction from large data.

## DISCUSSION AND CONCLUSION

Phylogenetic inference remains an indispensable framework for understanding biological relationships. At its core, phylogenetics attempts to faithfully describe the patterns of relatedness among biological entities. Beyond tracing evolutionary histories, phylogenetics allows researchers to organize biological diversity, identify species boundaries, track pathogen transmission, and interpret functional divergence at multiple biological scales (e.g., population-level diversity). As genomic data continue to expand exponentially, the demand for scalable, interpretable, and biologically faithful representations becomes increasingly essential (Deng et al., 2025; Piñeiro and Pichel, 2024).

Traditional phylogenetic workflows depend heavily on multiple sequence alignment, a computational and conceptual bottleneck that assumes positional homology across residues. However, this assumption often fails for highly divergent, recombined, or fragmented sequences, where alignments become ambiguous and error-prone (Chan and Ragan, 2013; Löytynoja et al., 2012). The growing field of alignment-free phylogenetics has sought to overcome this limitation by replacing residue-level comparisons with compositional or mathematical descriptors of sequence information (in a process called genetic encoding or representation). Coupled with the advances in machine learning, these approaches, now gaining increased attention (Chan et al., 2014), offer a more general framework for comparing biological sequences at scale.

Covary builds upon this paradigm, as did its predecessors, by introducing alignment-free inference in phylogenetics, with added translation-aware representation. However, unlike other models (reviewed somewhere else by Moeckel et al., 2024), its encoding architecture, derived from TIPs-VF, integrates codon boundary information and positional context within a unified vector representation, enabling the discrimination of mutational events that alter translational reading frames (De los Santos, 2025). This framework bridges compositional models with the biological relevance of translation, a dimension often ignored by these equally performing alignment-free methods (Fan et al., 2015; Zhang et al., 2019; Fan et al., 2022; Wen et al., 2014). The use of *k-mer*-based embeddings sensitizes Covary to perturbations and limitation previously documented of *k-mer*-based approaches (e.g., systematic errors), however, the improvements in *k-mer*-derived encoding in Covary also allows the extraction of meaningful relationship signals across sequences of variable length, reducing the dependency on explicit alignment or evolutionary assumptions encompassing translation-awareness. This tradeoff is believed to be more sensible than constraining the relationship estimation from models that often oversimplify or overestimate relatedness (Dimayacyac et al., 2023).

The results from Covary demonstrated strong discriminatory power in classifying both viral genome sequences and human cancer gene variants. Using complete SARS-CoV-2 genomes, Covary inferred major clusters among dominantly circulating variants, as previously demonstrated (Hozumi et al., 2021). Unlike conventional phylogenetic trees that reflect lineage-based temporal descent, topologies derived from Covary likely captures relationships shaped by genome-wide signatures and translational constraints. Previous experimental studies have shown that synonymous and codon-pair changes can alter SARS-CoV-2 fitness and host adaptation (Alonso and Diambra, 2020); hence, the produced clustering by Covary could be interpreted as evidence of composition-driven clustering rather than strict lineage separation. Although the implementation of codon-usage bias in the next iterations of Covary is in progress, future work should validate the hypothesis supporting composition-driven clustering by comparing codon usage and codon-pair bias between the observed clusters.

At the gene level, Covary effectively separated TP53 variants from GDC-TCGA datasets according to their translational consequence, distinguishing effectively between wild type, synonymous, missense, and frame-shifting mutations. This highlights the sensitivity of Covary to biologically meaningful perturbations that affect translation rather than mere sequence similarity. This observation primarily confirms the translation-aware capabilities of Covary. Moreover, clustering patterns within TP53 suggest possible functional partitioning related to truncating mutations, hotspot silent substitutions, or alternative isoform usage. Although these findings require further validation, they provide novel insights into how mutation-driven translational variation may influence p53 functional regulation and evolution (Kotler et al., 2018).

Covary demonstrated versatility across different biological resolutions, from gene-level mutation clustering to species- and genus-level phylogenetic reconstruction. When inferring the 18S rRNA gene tree among mammalian representatives, the topology produced by Covary agreed with that of traditional alignment-based methods like ETE3. This recapitulates accepted phylogenetic relationships without explicit MSA. Similarly, Covary accurately grouped bacterial genera and resolved subspecies clusters in *Lactobacillus*, underscoring the applicability of its translation-awareness in resolving even non-protein-coding sequences.

At the genomic scale, Covary successfully clustered viral and prokaryotic genomes into biologically interpretable groups, including taxonomically coherent families. The identification of inter-domain classification of bacterial and archaeal genomes in phylogenomic analysis illustrates the sensitivity of Covary to both perturbations in genome complexity and flexibility to reveal patterns shaped by horizontal gene transfer, compositional convergence, or variable evolutionary rates. However, such observation also suggests that the success of similar models are inherently applicable to Covary, as have been presented in this report. Hence, the success of Covary was not due to random embeddings or by chance.

The results presented collectively indicate that Covary not only performs comparably to conventional model-based tools but may also capture broader dimensions of genomic similarities beyond alignment-defined homology. Although only 10 prokaryotic chromosomal genomes were analyzed in this report, using modest computational resources, Covary can be easily scaled to process higher sample size when performed on advanced computing infrastructure. Nonetheless, work is already in progress to scale this capacity to hundreds or thousands of genomes in the coming years.

Despite that Covary was tested on Google Colab using modest computational resource (free tier), it already managed to analyze almost a thousand (n=906) complete SARS-CoV-2 genomes in minutes. This demonstrates its capacity for immediate deployment for viral epidemiological studies and flexibility for larger-scale phylogenomic analyses when subsumed on more powerful systems. Its alignment-free and translation-aware design allows it to analyze diverse sequence datasets efficiently, making it suitable for biocomputational and comparative genomics applications where rapid and scalable relationship estimation is needed. The ability of Covary to resolve taxonomic structure and detect fine-scale genomic variation also supports species identification, classification, and biodiversity monitoring, thereby extending its relevance to taxonomic and ecological studies. Moreover, by modeling relationships directly from codon-bound sequence embeddings, Covary enables researchers to explore evolutionary patterns and functional divergence without being constrained by traditional alignment models, positioning it as a unifying analytical tool for next-generation phylogenetic and genomic research.

The broader vision of Covary aligns with the next frontier in molecular systematics, which is whole-genome phylogenetics. Traditional substitution models, parsimony methods, and likelihood-based frameworks, while foundational, are increasingly challenged by the sheer volume and diversity of available genomic data, including model limitations. The future of phylogenetics will rely on algorithms capable of integrating information from entire genomic spaces into coherent, interpretable relationship networks (Piñeiro and Pichel, 2024; Fan et al., 2015). The translation-aware, *k-mer*-based embeddings of Covary represent an early and crucial step toward this future, enabling rapid and objective estimation of biological relationships at genome scale.

Despite its promise, Covary presents several limitations that guide future refinements. First, while Covary tolerates variable-length sequences, it was found that excessive gaps (>75% relative to informative sites) can distort inferred relationships (data not shown). Users should ensure minimal missing regions or pre-trim poorly covered sequences. Second, because Covary uses *k-mer* representations rather than alignment-based correction, erroneously gapped regions that disrupt codon frames can introduce artificial dissimilarities. Manual inspection or preprocessing is recommended for such cases. Third, the default 6-mer encoding enhances translation awareness but makes the model sensitive to positional perturbations. The next iterations of Covary encoder may explore 3-mer optimization for improved efficiency and interpretable branch-length estimation. Fourth, distance metrics in Covary are derived from vector-space dissimilarities rather than evolutionary rate estimations. Thus, current branch lengths reflect relational magnitude rather than time. Future model calibration could incorporate *k-mer* frequency vocabularies to approximate substitution distances. Fifth, although Covary captures codon boundaries, it still treats sequences as symbolic composites of *k-mers* rather than as full derivatives of the genetic code. Consequently, analyses requiring explicit codon substitution modeling remain partially outside its scope, but not impossible. Sixth, present workflows handle large datasets in memory, limiting scalability to thousands of genomes. Parallelization, chunked processing, and GPU optimization are planned to expand the performance of Covary in large-scale phylogenomics. Seventh, Covary has yet to be systematically benchmarked with commonly used models, such as likelihood or Bayesian frameworks, and molecular markers. However, it is hypothesized that these classical models will eventually yield to data-driven, high-dimensional representations capable of processing entire genomes, an arena where Covary and other machine learning-based models are already positioned to dominate. Lastly, tree search in Covary is done manually. Work to automate best-fitting tree is already in progress for the next iterations or updates on Covary.

Covary represents a step toward a new generation of relationship-estimation frameworks that are translation-aware, alignment-free, and inherently scalable. Its demonstrated versatility across mutation-level, gene-level, and whole-genome analyses underscores its potential to unify phylogenetic inference under a single, data-driven system. While refinements remain necessary, particularly in optimizing scalability and interpretability, Covary and other large data methods lay the groundwork for the future of genome-scale relationship modeling. As genome sequencing becomes universal and cheaper, tools like Covary will enable a more faithful, relevant, and objective understanding of life’s relationships, fulfilling a long-standing goal of molecular phylogenetics, that is to represent biological ‘relatedness’ in its most comprehensive form.

