## Supplementary Figure for "Covary: A translation-aware framework for alignment-free phylogenetics using machine learning"

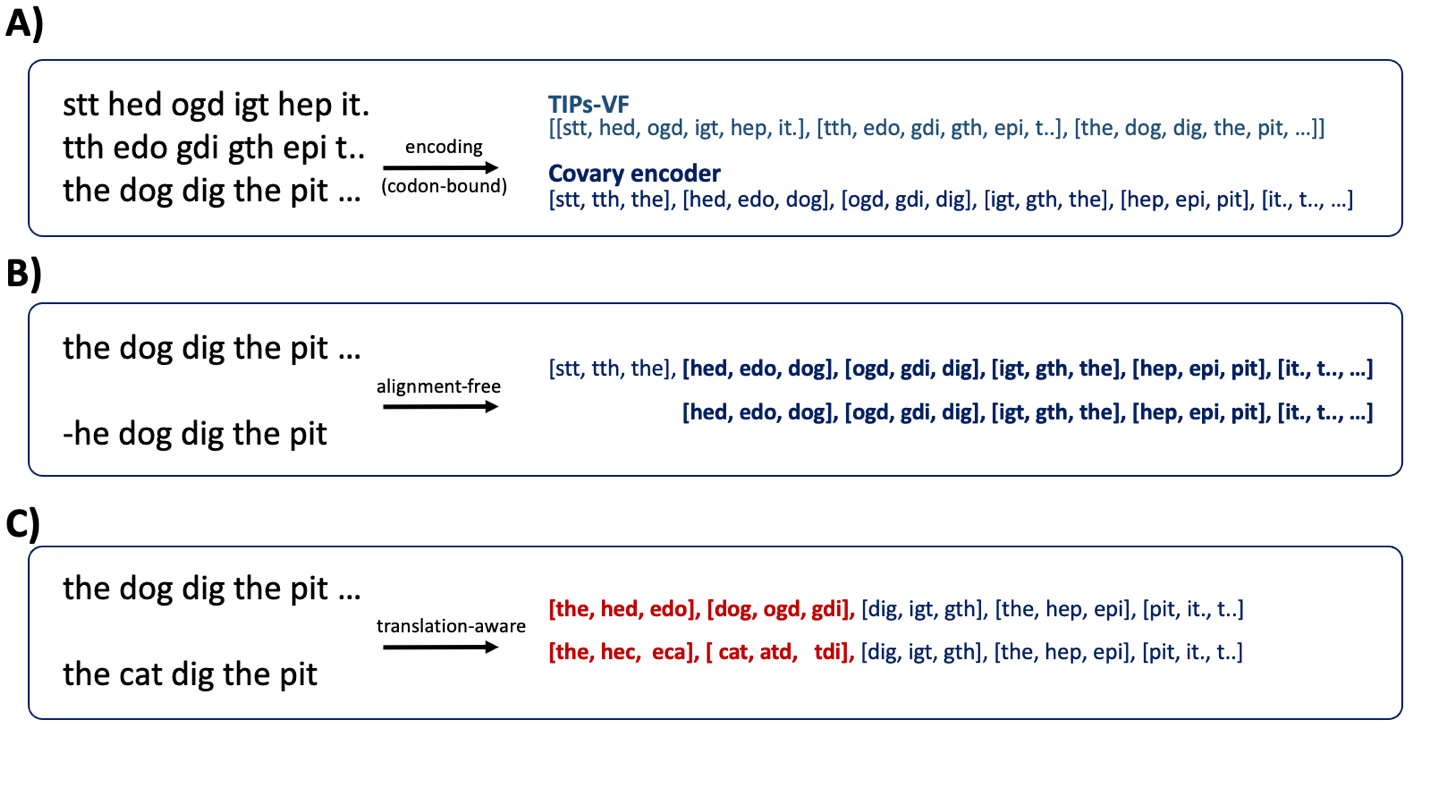

**Supplementary Figure 1. The Covary encoder module illustrating alignment-free and translation-aware features.** A) Comparative schematic of the genetic encoding processes implemented in TIPs-VF and the Covary encoder. B) Conceptual representation of the alignment-free encoding strategy in Covary. C) Illustration of translation-aware encoding approach, showing the integration of codon boundary information across reading frames using the Covary encoder. Legend: The minus (–) symbol in (B) indicates sequence truncation.

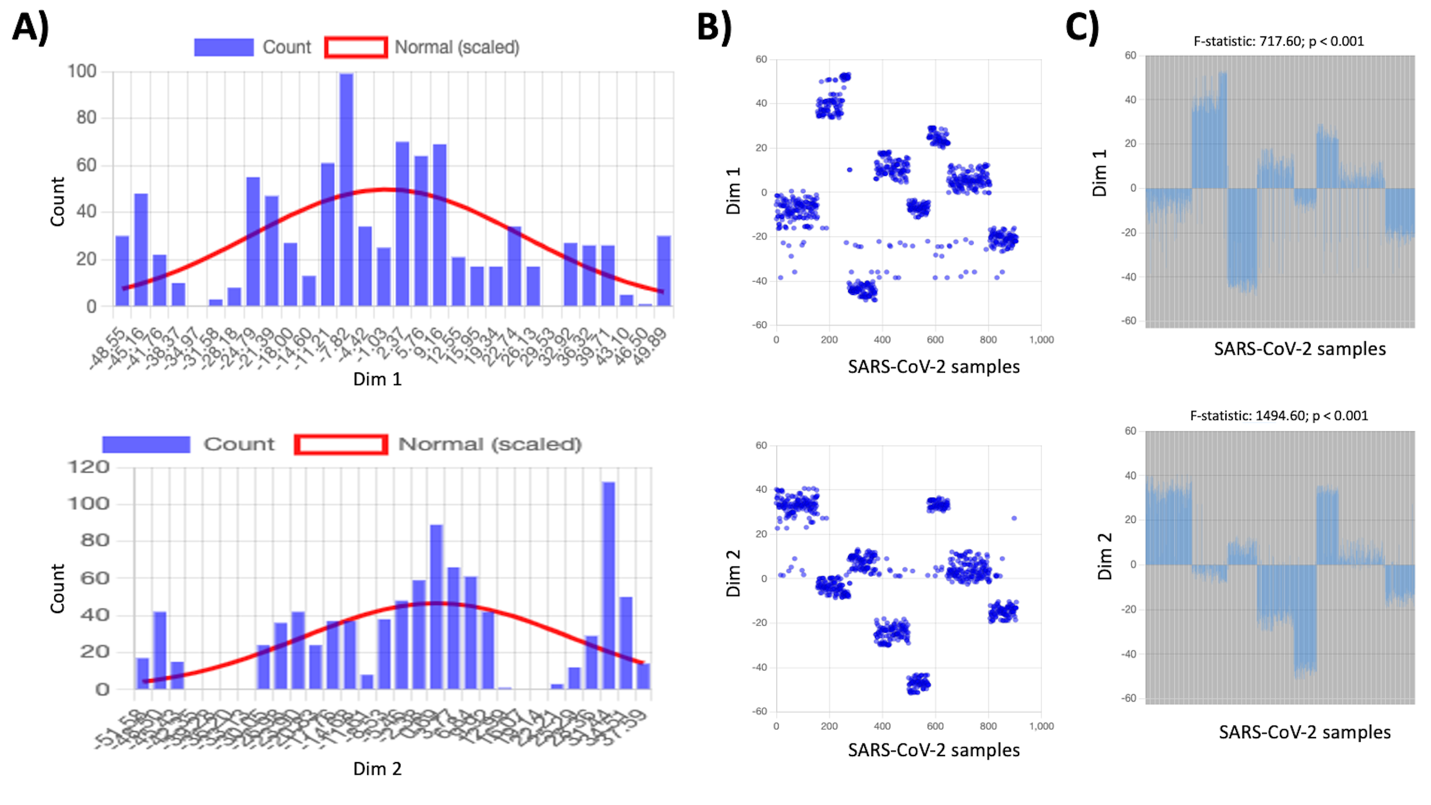

**Supplementary Figure 2. Statistical validation of Covary-derived distance matrices using SARS-CoV-2 whole-genome embeddings.** A) Histogram of normal distribution analysis for Dimensions 1 and 2 (Dim 1 and Dim 2). B) Scatter plot and (C) bar graph visualizations of Dim 1 (upper) and Dim 2 (lower) embeddings derived from SARS-CoV-2 genome sequences. Cluster differences were assessed using one-way ANOVA (p < 0.0001; extremely significant). Statistical computations were performed using PlethoCalc.

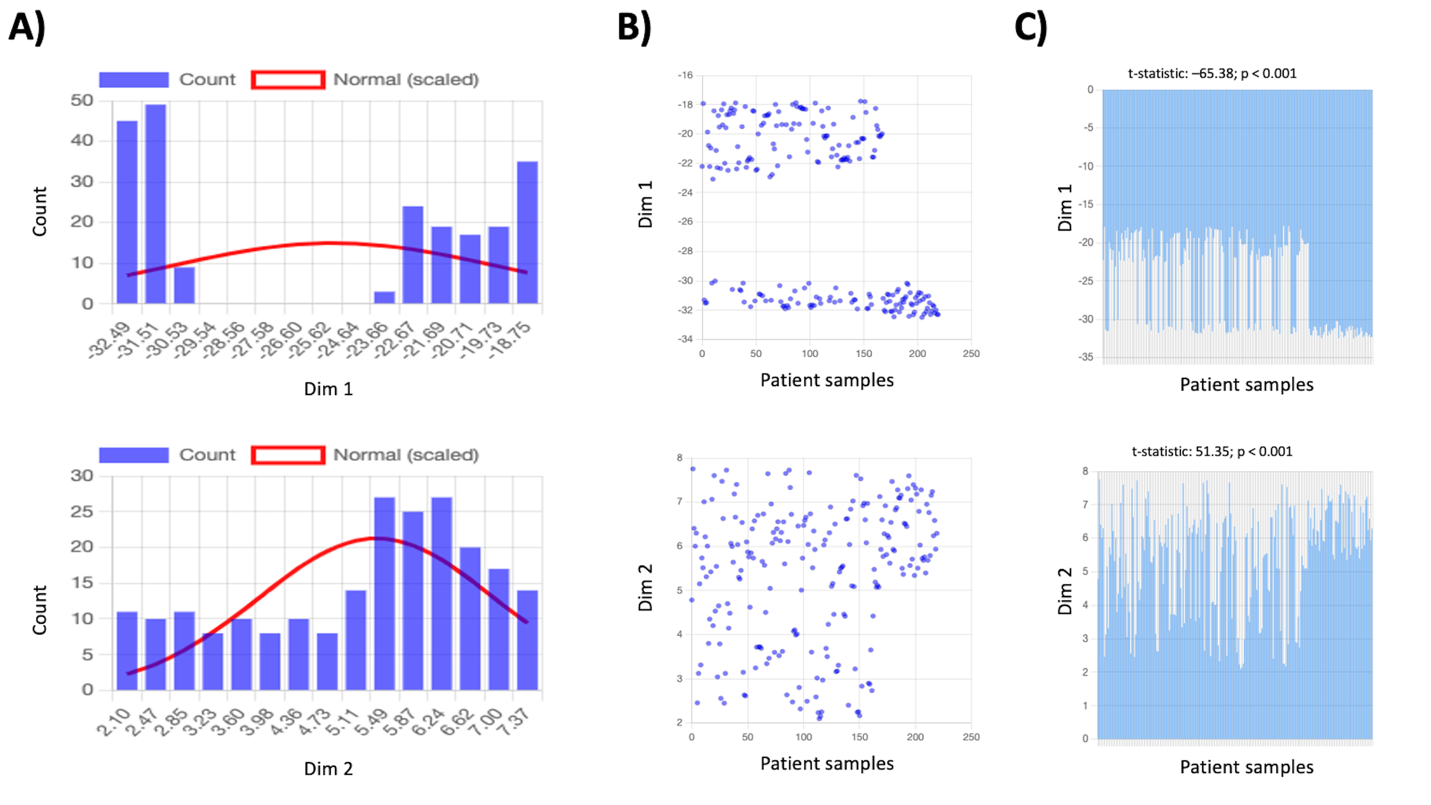

**Supplementary Figure 3. Statistical validation of Covary-derived distance matrices using TP53 mutational profiles from cancer patients.** A) Histogram of normal distribution analysis for Dimensions 1 and 2 (Dim 1 and Dim 2). B) Scatter plot and (C) bar graph visualizations of Dim 1 (upper) and Dim 2 (lower) embeddings derived from patient-specific TP53 sequences. Cluster differences were evaluated using a one-sample t-test (p < 0.0001; extremely significant). Statistical analyses were conducted using PlethoCalc.

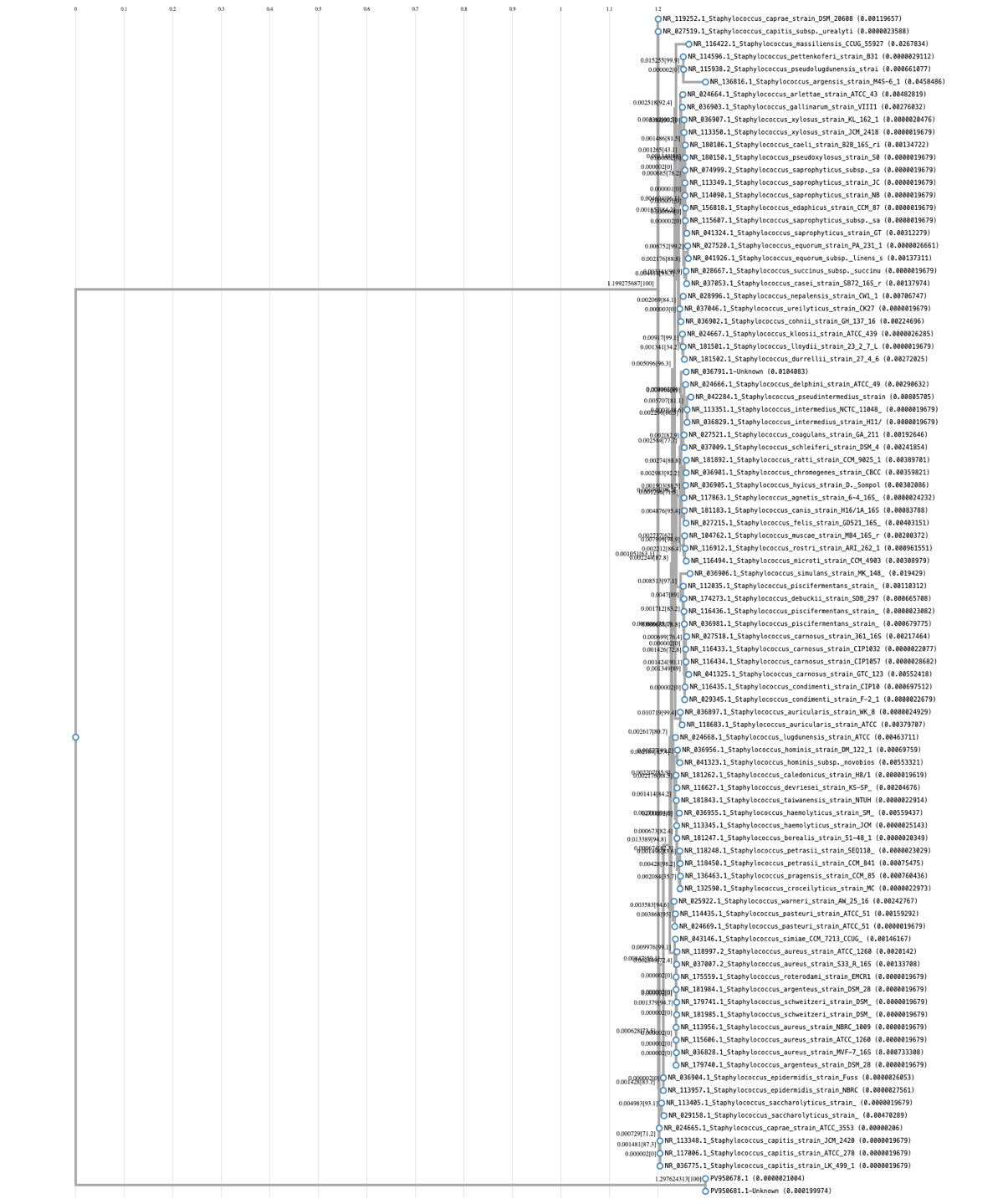

**Supplementary Figure 4. Species identification and outgroup detection using 16S rRNA markers with ETE3.** Phylogenetic reconstruction accurately placed both outgroup and ingroup sequences into their respective clades. The taxonomy-verified Staphylococcus sequence (NR_036791.1) was correctly positioned within the ingroup, while the SARS-CoV-2 genome (PV95068.1) was appropriately assigned as an outgroup, confirming correct taxonomic placement by ETE3.

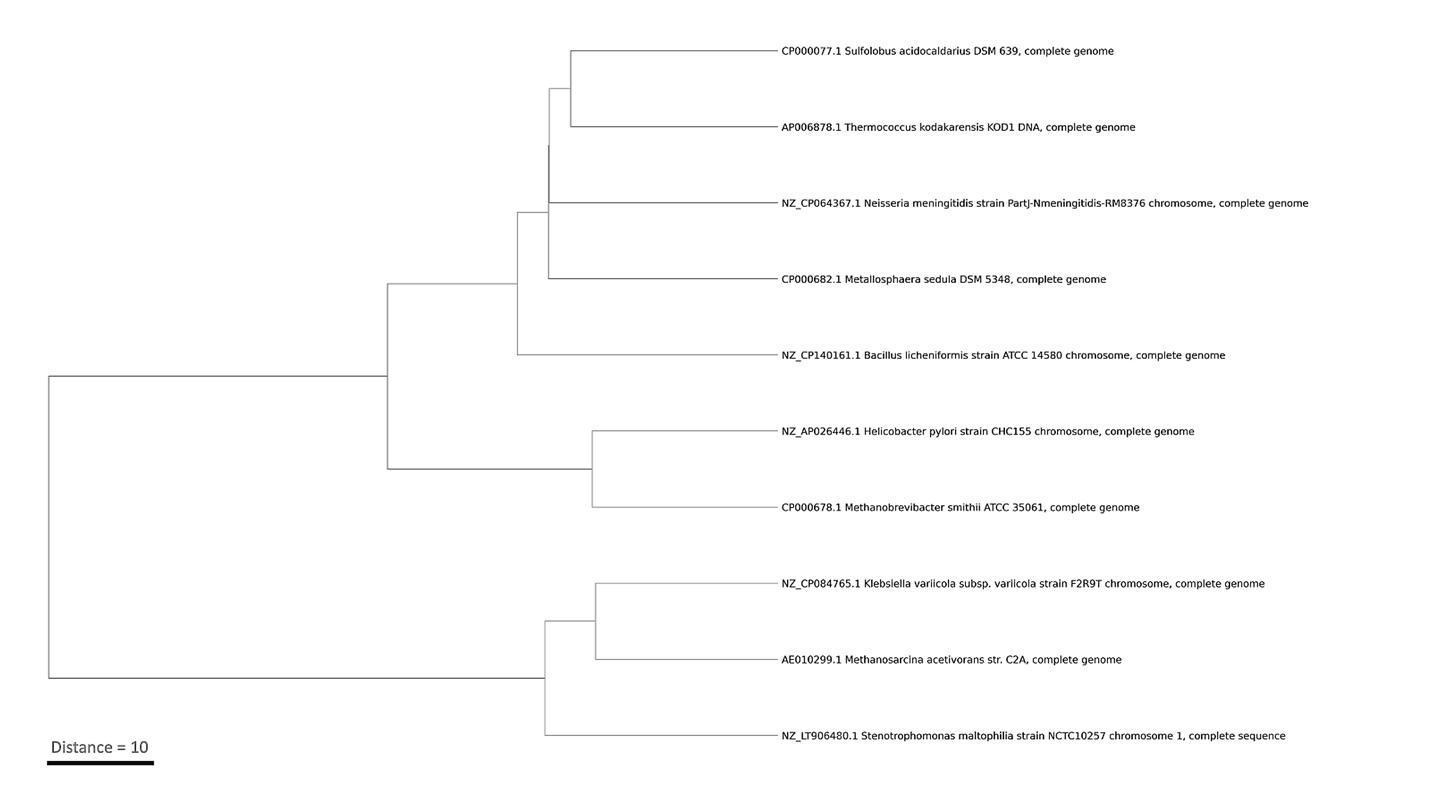

**Supplementary Figure 5. Whole-genome phylogenomic analysis of archaeal and bacterial species using Covary.** Genome-scale analysis with Covary resolved two major clades encompassing archaeal and bacterial taxa while revealing unexpected inter-domain relationships. Archaeal species such as *Methanosarcina acetivorans, Methanobrevibacter smithii, Metallosphaera sedula, Thermococcus kodakarensis*, and *Sulfolobus acidocaldarius* clustered closely with bacterial taxa, suggesting that Covary can detect genomic signatures indicative of horizontal gene transfer, compositional convergence, or shared evolutionary constraints. This result also confirms the associated perturbations and incongruencies associated with *k-mer*-derived phylogenies.
